# Fluctuation-Based Super-Resolution Traction Force Microscopy

**DOI:** 10.1101/772947

**Authors:** Aki Stubb, Romain F. Laine, Camilo Guzmán, Ricardo Henriques, Guillaume Jacquemet, Johanna Ivaska

## Abstract

Cellular mechanics play a crucial role in tissue morphogenesis and homeostasis and are often misregulated in disease. Traction force microscopy (TFM) is one of the key methods that has enabled researchers to study fundamental aspects of mechanobiology; however, the power of TFM is limited by poor resolution and low throughput. Here, we propose a simplified protocol and imaging strategy, relying on super-resolution microscopy enabled by fluorophore fluctuation analysis, to enhance the output of TFM, by increasing both bead density as well as the accuracy of bead tracking in TFM gels. Our analysis pipeline can be used on either camera-based confocal or widefield microscopes and is fully compatible with available TFM analysis software. In addition, we demonstrate that our workflow can be used to gain biologically relevant information and is suitable for long-term live measurement of traction forces even in light-sensitive cells. Finally, we propose that our strategy could be used to considerably simplify the implementation of TFM screens. Our streamlined protocol can be performed with minimal hardware and software investment, and has the potential to standardize high-resolution TFM.

## Main text

Cell adhesion to the extracellular matrix (ECM) is a fundamental feature of multicellular life and it is finely tuned during virtually every cellular process including cell migration, cell proliferation and cell fate. Cells do not only passively attach to the ECM but also constantly apply forces on ECM molecules and actively remodel their microenvironment (Conway and Jacquemet, 2019). The major cellular structures responsible for transmitting these forces to the ECM are focal adhesions (Conway and Jacquemet, 2019). These multiprotein signaling platforms also translate physical forces into intracellular biochemical signaling cascades (Kechagia et al., 2019). The ability of cells to apply mechanical forces on their environment is emerging as one of the key regulators of tissue patterning and morphogenesis (Heisenberg and Bellaïche, 2013; Wickström and Niessen, 2018), while dysregulation of this process is associated with diseases including aging, fibrosis and cancer (Duscher et al., 2014; Segel et al., 2019; Broders-Bondon et al., 2018). As our understanding of mechanobiology is rapidly unveiling promising novel therapeutic opportunities (Lampi and Reinhart-King, 2018), the development of methods that can facilitate their study and make them more widely available is of paramount importance.

While several strategies can be used to map and quantify the forces exerted by cells on their microenvironments, traction force microscopy (TFM) is one of the most convenient and widely used methods (Roca-Cusachs et al., 2017). To perform TFM, cells are allowed to adhere to a deformable material of defined stiffness, classically a polyacrylamide (PAA) gel, containing fluorescent beads. The forces exerted by cells on their substrate can then be monitored as a function of bead movement within the gel. As the stiffness and the elastic modulus of the gel are pre-established, bead displacement can then, using mathematical equations, be converted into a read-out of local forces (Style et al., 2014).

The sensitivity and accuracy of TFM is directly linked to the ability to detect and track moving beads within a thick gel, and is generally limited to the detection of forces at the micron-scale (Roca-Cusachs et al., 2017; Style et al., 2014). As focal adhesions, the cellular structures involved in force generation and ECM remodelling, can be much smaller (Changede et al., 2019), there is a need to develop methods that can map cellular forces at high spatial resolution. For TFM, this can be achieved by (1) increasing the number of trackable beads (Plotnikov et al., 2014; Colin-York et al., 2016, 2019) and by (2) improving the computational algorithms used to map cellular forces (Han et al., 2015). Multiple imaging and sample preparation strategies have been developed to increase the number of trackable beads in TFM experiments, each with unique strengths and shortcomings (Table 1) (Colin-York et al., 2016, 2019; Plotnikov et al., 2014), while improvements in TFM algorithms often require further biological assumptions (Zündel et al., 2017) as well as heavy computational processing power (Han et al., 2015). Here we propose a simplified protocol that can resolve densely packed beads, using software enabling super-resolution microscopy through fluorophore intensity fluctuation analysis. For simplicity, these algorithms are hereafter termed fluctuation-based super-resolution (FBSR) imaging. In this study, we demonstrate that FBSR combined with TFM considerably enhances traction force outputs.

**Table 1.** High resolution TFM of cells plated in 2D.

|  | **STED TFM**<br>(Colin-York et al., 2016) | **SIM TFM**<br>(Colin-York et al., 2019) | **High-resolution TFM**<br>(Plotnikov et al., 2014) | **FBSR TFM**<br>(This study) |
| --- | --- | --- | --- | --- |
| **Required microscope** | STED equipped with long working distance objective | SIM | Confocal | Camera-based confocal or widefield microscope |
| **Live-cell imaging friendly** | No | No | Yes | Yes |
| **Number of other fluorescent channels typically available** | 1 | 3 | 2 | 3 |
| **Expected bead density** | 2.2 beads per $\mu\text{m}^2$ | 1.0 beads per $\mu\text{m}^2$ | Information not available | 1.2 beads per $\mu\text{m}^2$ |
| **TFM analysis** | Classical analysis pipeline can be applied. Open access software readily available. | Normal pipeline for 2D, specific for 3D. | Specific analysis pipeline required. No specific software readily available. | Classical analysis pipeline can be applied. Open access software readily. |
| **Control for imaging artefacts** | No | No | Not required | Yes |
| **Bead size used** | 40nm | 100nm | 40nm | 40nm |
| **Bead tracking** | 2D tracking | 3D tracking using dedicated software | 2D tracking | 2D tracking |

FBSR harnesses the intrinsic property of fluorophores that, when excited with continuous light, display random variation in intensity over time due to transitions between fluorescent and non-fluorescent states (Bagshaw and Cherny, 2006; Linde and Sauer, 2014). After capturing these intensity oscillations (typically tens to hundreds of images), algorithms such as Super-resolution Optical Fluctuation Imaging (SOFI, (Dertinger et al., 2009)), Super-Resolution Radial Fluctuations (SRRF, (Gustafsson et al., 2016)) or autocorrelation two-step deconvolution (SACD) (Zhao et al., 2018) can be used to predict the location of fluorophores at improved resolution. Here, we demonstrate that applying FBSR imaging to TFM can substantially improve the resolution of traction force measurements. Importantly, our strategy only requires off-the-shelf reagents and access to commonly available widefield or camera-based confocal microscopes. Our analysis pipeline is fully compatible with freely available TFM analysis software. Compared to other superresolution (SR) modalities, FBSR minimises phototoxicity and in our protocol is suitable for long-term live measurement of traction forces even in light-sensitive cells. With our detailed protocol on implementing fluctuation-based TFM, we believe that the cell biology community can rapidly adopt this method to investigate mechanobiology.

### Increasing the bead density of TFM gels using fluctuation-based microscopy

To increase the number of trackable beads in our TFM experiments, we decided to employ FBSR (Figure 1a) as it has several advantages over other SR modalities (Table 1). Of note, FBSR is easy to implement, is compatible with most pre-existing microscopes including spinning-disk confocal (SDCM) and widefield systems (Gustafsson et al., 2016). and is only mildly phototoxic with improved resolution being achieved using illumination intensities typical for conventional fluorescence imaging (Culley et al., 2018b).

**Fig. 1.**
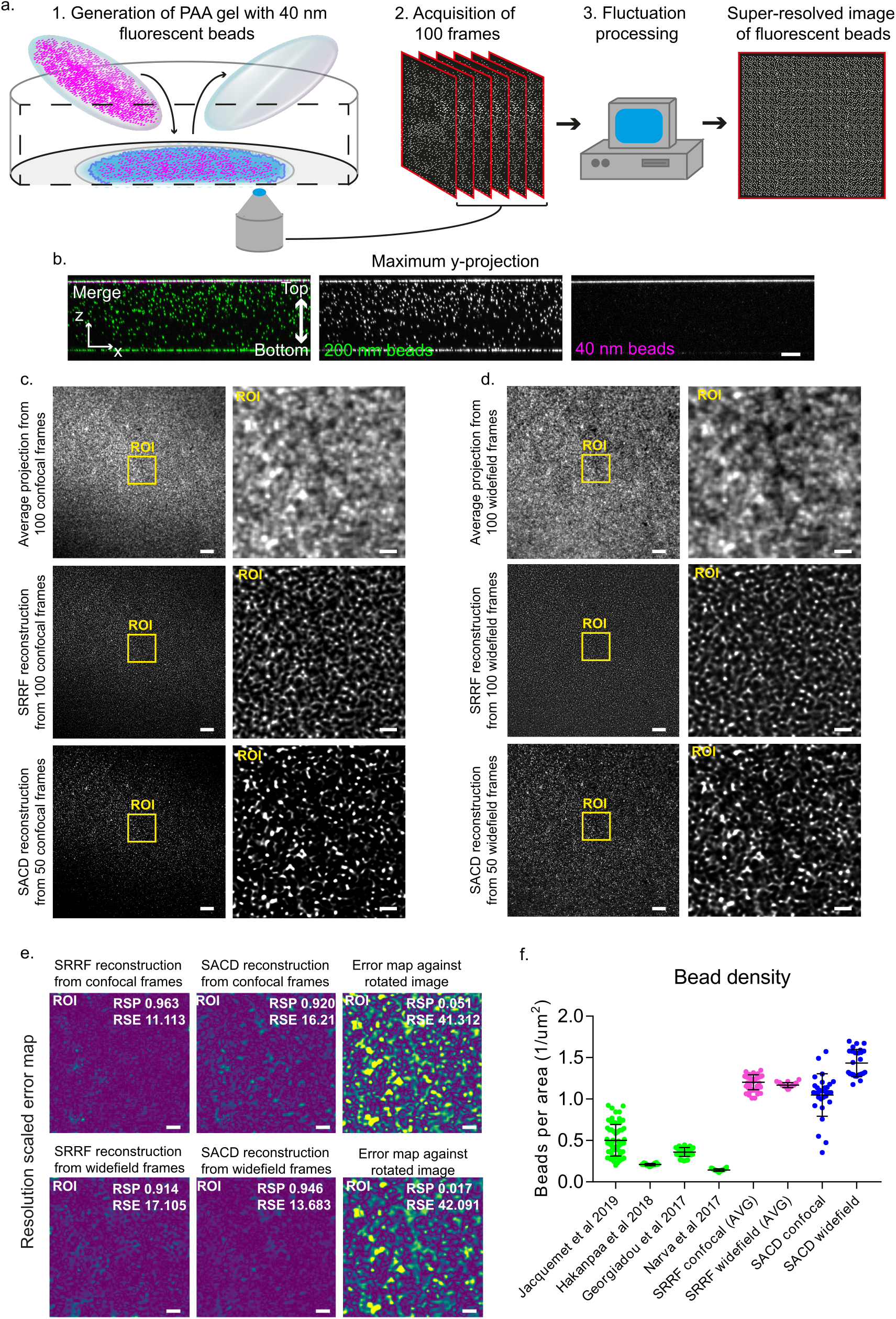
FBSR processing enhances bead recognition in TFM specific PAA gels. (**a**) Cartoon illustrating the key steps used to produce TFM gels for FBSR imaging. 1) A TFM gel were 40 nm fluorescent beads are embedded only on the topmost layer of the gel is generated. 2) TFM gels are then imaged using a camera-based confocal or widefield microscope. To allow for FBSR processing, each field of view is imaged 100 times. 3) SR images are then generated using available FBSR algorithms such as LiveSRRF or SACD. (**b**) TFM gel containing both 200 nm beads (distributed throughout the gel, classic protocol) and 40 nm beads (distributed only at the top of the gel, new protocol described here) were imaged using SDCM. A maximal horizontal projection is displayed to highlight bead distribution. Scale bar 10 µm. (**c-d**) A TFM gel prepared using our improved protocol (40 nm beads embedded only at the top) was imaged using SDCM (**c**) or widefield (**d**) modes. To allow for FBSR, 100 frames were recorded. Average projections as well as LiveSRRF and SACD processed images are displayed. For each condition, the yellow square highlights a region of interest (ROI) that is magnified. All images are from the same field of view. Scale bar: (main) 10 µm; (inset) 2 µm. (**e**) Resolution scaled error maps of the LiveSRRF and SACD images displayed in c and d (ROI only) generated with NanoJ-SQUIRREL. Maps are color-coded to visualize areas of low (purple) and high error (yellow). As a control, the same analysis was performed using a reference frame that was rotated 90°. Scale bars 2 µm. (**f**) Graph showing bead densities (beads per square micrometre) measured from multiple published TFM datasets (Jacquemet et al., 2019; Hakanpaa et al., 2018; Georgiadou et al., 2017; Närvä et al., 2017) and from TFM gels (improved protocol described here) imaged using either SDCM or widefield followed by FBSR processing using LiveSRRF or SACD.

Classically, when performing TFM experiments, 200 nm fluorescent beads are embedded throughout the PAA gel (Style et al., 2014). This can result in substantial out-of-focus light, which limits the use of widefield microscopes for TFM. To increase the number of trackable beads in our TFM experiments and enable better quantification of cellular forces, we used gels containing densely packed 40 nm fluorescent beads. In addition, to optimize TFM gels for FBSR, we developed a simplified gel casting protocol where the 40 nm beads are embedded only on the topmost layer of the gel (Figure 1b and Supplementary Figure 1a). This is achieved by pre-coating the top coverslip, used to flatten the gel solution prior to casting, with the beads instead of mixing the beads within the gel solution itself (Supplementary Figure 1a). Importantly, using the FBSR algorithms LiveSRRF and SACD and our optimized protocol, we were able to resolve the 40 nm beads located on top of the TFM gel using both SDCM and widefield microscopes (Figure 1c and 1d). To ensure that as few artefacts as possible were introduced during the FBSR reconstruction process, the image quality was assessed using NanoJ SQUIRREL (Culley et al., 2018a) and the Resolution Scaled Pearson’s correlation (RSP) and Resolution Scaled Error (RSE) parameters were calculated by the software (Figure 1e). In addition to these parameters, when choosing the reconstruction settings, the amount of beads detected and the absence of patterning in the final image were also taken into consideration (Supplementary Figure 1b and 1c).

Prior to FBSR, our confocal-based TFM analyses have yielded between 0.2 to 0.5 trackable beads per square micrometre (Jacquemet et al., 2019; Närvä et al., 2017; Georgiadou et al., 2017; Hakanpaa et al., 2018) (Figure 1f), in agreement with values reported by others (Colin-York et al., 2016). Here, by implementing FBSR, and conservative reconstruction parameters, we were able to substantially increase the number of trackable beads to 1.2 beads per square micrometre (Figure 1f). This is a modest improvement over a protocol using structured illumination microscopy (Colin-York et al., 2019) (1 bead per square micrometre) but remains inferior to another protocol based on STED imaging within small fields of view (2.2 beads per square micrometre) (Table 1) (Colin-York et al., 2016). Interestingly, FBSR performed especially well when images were acquired using widefield microscopy as the final SR images were more homogenous (Figure 1d). When the images were acquired using SDCM, the corners of the field of view were often off focus due to uneven / wrinkled gels resulting in much lower beads density in these areas. In particular SDCM images reconstructed by SACD appear to be especially sensitive to out of focus light, arisen from uneven gels, leading to more variable beads density than the other imaging modalities (Figure 1c).

### Implementation of fluctuation-based traction force microscopy

In a typical TFM experiment, beads are imaged before (Pre) and after (Post) removing cells, the Pre and Post images are then aligned and the beads are detected and then tracked in both images (Style et al., 2014). From the tracking data, bead displacement maps and force maps can be generated using available TFM software (Han et al., 2015; Tseng et al., 2012). FBSR-TFM follows the same workflow with the addition of image reconstruction prior to the Pre and Post image alignment (Figure 2a).

**Fig. 2.**
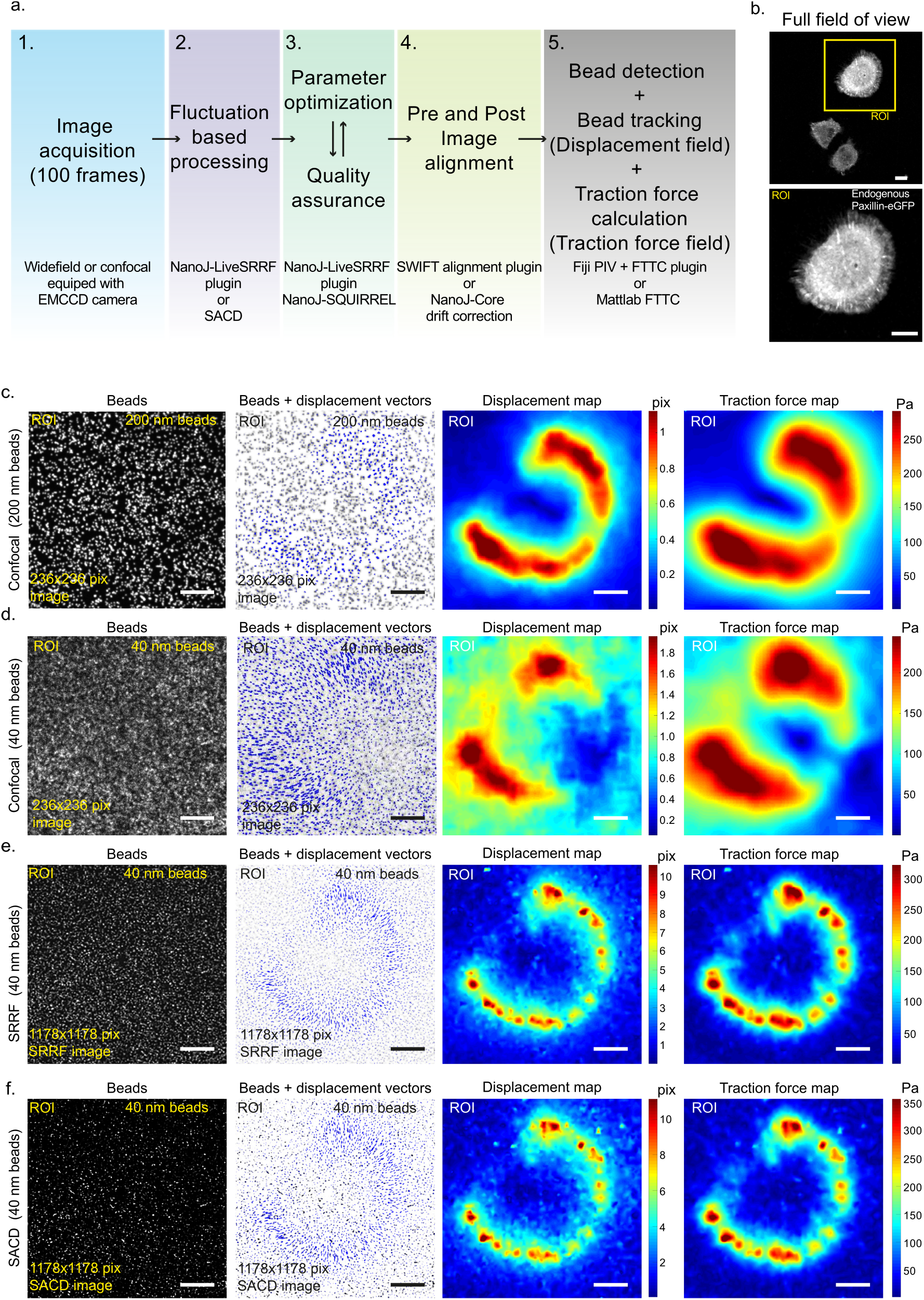
Implementation of FBSR to TFM. (**a**) Schematic pipeline of a TFM experiment that includes FBSR and image quality control. The software needed to complete each step are listed. (**b-f**) To assess the improvement generated by FBSR-TFM over the classically used confocal-based TFM, U2OS cells expressing endogenously tagged paxillin (**b**) were plated on 9.6 kPa gels containing both 40 nm and 200 nm beads (as in Figure 1b) and TFM analyses were performed (as in panel **a**) on the ROI (yellow square, **b**). SDCM images of 200 nm (**c**) and 40 nm beads (**d**) and FBSR images of the 40 nm beads (LiveSRRF (**e**); SACD (**f**)) were used for TFM analysis using a MATLAB-based software (Han et al., 2015). For each method, images of beads alone and beads (black) + displacement vectors (purple arrows, length scaled up by 2) and maps of bead displacement and traction force are displayed. The magnitudes of bead displacement and traction force are color-coded as indicated. Scale bars 10 µm. Analyses of the full field of view from panel b can be found in supplementary Figure 2.

To assess the improvement generated by FBSR-TFM over classically used confocal-based TFM, cells were plated on gels containing both 200 nm beads (distributed throughout the gel, classic protocol) and 40 nm beads (distributed only at the top of the gel, new protocol described here) (Figure 1b, Figure 2b-c and Supplementary Figure 2a-b). This strategy enabled us to measure and visualise traction forces using both methodologies within the same field of view (Figure 2c-f). Using SDCM imaging of the 200 nm beads (classic confocal TFM) or of the 40 nm beads, we were able to track 8253 beads and 11328 beads, respectively (full field of view, Supplementary Figure 2a-b). In contrast, FBSR imaging of the 40 nm beads, yielded 20799 (LiveSRRF processing using 100 frames) and 22908 trackable beads (SACD processing using 50 frames) within the same field of view. The SDCM-based TFM generated displacement and traction force maps that closely recapitulated the shape of the cell (Figure 2b-f). However, at this resolution, areas corresponding to cell-ECM contacts such as focal adhesions could not be pinpointed and results were not substantially improved when using the SDCM images of the 40 nm beads (Figure 2d). Strikingly, when applying the same TFM pipeline to the FBSR images of the 40 nm beads (regardless of the FBSR method used), drastically improved the resolution of the displacement and force maps. In particular, defined regions of high force were specifically detected at the cell perimeter, which could correspond to focal adhesions (Figure 2e and 2f). In addition, due to enhanced bead tracking, we could better segregate cellular regions corresponding to weaker forces. While the final force maps are affected by the algorithm / mathematical framework used to perform force reconstruction (Han et al., 2015), the bead displacement maps are a direct reflection of the amount of beads used as well as the quality of bead tracking (Roca-Cusachs et al., 2017; Plotnikov et al., 2014). In particular, errors in displacement measurements caused by a lack of accuracy in the tracking routines strongly affect the resolution of TFM. Notably, in the case of the SDCM TFM, large beads are only tracked over subpixel movements, which is likely to lead to tracking inaccuracies (figure 2c-f). In contrast, in the case of FBSR-TFM the tracking accuracy is likely to be improved as 1) the beads are smaller and 2) they are now tracked over several pixels (due to the smaller pixel size of the FBSR images). Overall we believe that FBSR improves the TFM outputs by both increasing the bead density (more data points) and by refining the accuracy of bead tracking.

### Versatility of fluctuation-based TFM

One of the advantages of FBSR is that it can easily accommodate multicolor imaging. To demonstrate this capability, we set out to measure forces in cells endogenously tagged for paxillin, a marker of focal adhesions. In this case, FBSR not only enhanced bead tracking and identification but also the resolution of paxillin-positive focal adhesions (Figure 3a, Supplementary Figure 3a). Importantly, this multicolor imaging capability combined with enhanced image quality, enabled us to confirm that the observed regions of higher force correlate with the localisation of cell-ECM adhesions (Figure 3a and 3b).

**Fig. 3.**
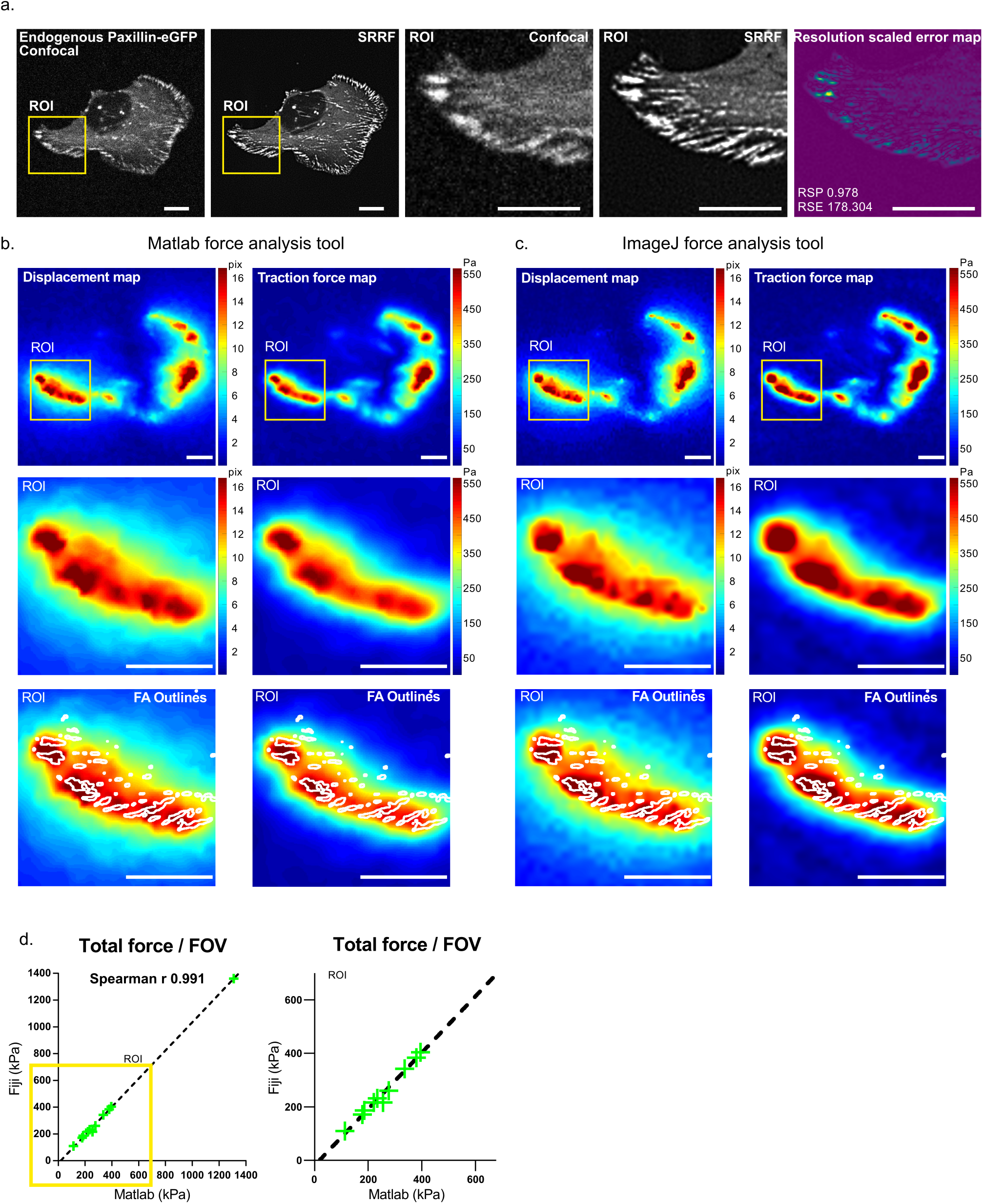
Applying FBSR-TFM to cell biological experiments. (**a-d**) U2OS cells expressing endogenously tagged paxillin were plated on 9.6 kPa gels containing 40 nm beads and were imaged using an SDCM. In this dataset, both the beads and paxillin were imaged for FBSR processing. (**a**) Representative image of paxillin-positive focal adhesions before and after FBSR processing using LiveSRRF. Yellow squares highlight a ROI that is magnified. For the ROI, the resolution scaled error map is also displayed as in Figure 1e. Scale bars 10 µm. (**b,c**) TFM analyses were performed on the FBSR images of the 40 nm beads displayed in figure 3a using a MATLAB-based software (**b**) or using the ImageJ PIV and FTTC plugins (**c**). Representative bead displacement and traction force maps generated by these two software are displayed as described previously. The yellow squares highlight the same region of interest that is magnified. On the bottom panel, the outlines of the LiveSRRF enhanced focal adhesions (figure 3a) are overlaid (white) with the bead displacement and traction force maps. (**d**) Correlation between the force measurements performed with either the MATLAB or the ImageJ TFM pipeline (n = 11 fields of view). Correlation coefficient r measured using the Spearman’s Rank-Order. Analyses of the full field of view for the representative image can be found in Supplementary Figure 3.

To demonstrate that FBSR is compatible with multiple existing TFM pipelines, we compared the displacement and force maps generated by two freely available software (MATLAB (Han et al., 2015) or ImageJ-based software (Tseng et al., 2012); see methods for details). Regardless of the software used, the displacement and force maps matched well to the outline of the focal adhesions (Figure 3b,c, Supplementary Figure 3b,c). Furthermore, the total amount of forces measured using these two software showed a remarkable correlation (spearman r 0.991) (Figure 3d). It is noteworthy that the MATLAB-based software produced more well-defined bead displacement and force maps compared to the ImageJ-based software. This is likely due to the implementation of a high-resolution bead-tracking algorithm in the MATLAB software compared to PIV in ImageJ (Han et al., 2015; Tseng et al., 2012).

### Biologically relevant applications of fluctuation-based TFM

Next, we sought to demonstrate that our improved TFM pipeline could be used to answer biologically relevant questions. In particular, TFM is very commonly used to assess how a protein or a drug treatment influences the ability of cells to exert forces on their environment (Jacquemet et al., 2019; Hakanpaa et al., 2018; Närvä et al., 2017). For this purpose, we aimed to cause a mild perturbation to simulate a plausible biological response and treated cells with either DMSO or the myosin II inhibitor blebbistatin for 15 minutes (Figure 4a and 4b). FBSR imaging of focal adhesions and FBSR TFM measurements were performed before and after the treatments to allow the quantification of force changes in each cell. Importantly, 15 minutes treatment with blebbistatin (at the concentration used) was not sufficient to trigger the collapse of focal adhesions (Figure 4b). While blebbistatin treatment triggered a notable decrease in the traction forces exerted by cells (Figure 4a,b), this effect was masked by the high variability in both DMSO and blebbistatin-treated cell populations when comparing the 15-minute time point only (Figure 4c). Therefore, we directly compared cells before (time point 0) and after treatments (time point 15 min) and calculated the fold change in traction force and strain energy (SE) for each condition (DMSO and blebbistatin). We found that blebbistatin treatment significantly decreased both traction forces and SE (Figure 4d), in line with our traction force maps. These results indicate that the negative effect of blebbistatin on cellular forces, usually associated with a dramatic loss in focal adhesions, occurs, and can be detected by FBSR-TFM, at earlier stages preceding focal adhesion disassembly. In addition, our FBSR-TFM analysis pipeline highlights the value of performing TFM prior to and after a perturbation on the same cell to remove cell-to-cell variability and thus to more accurately detect changes in cellular forces.

**Fig. 4.**
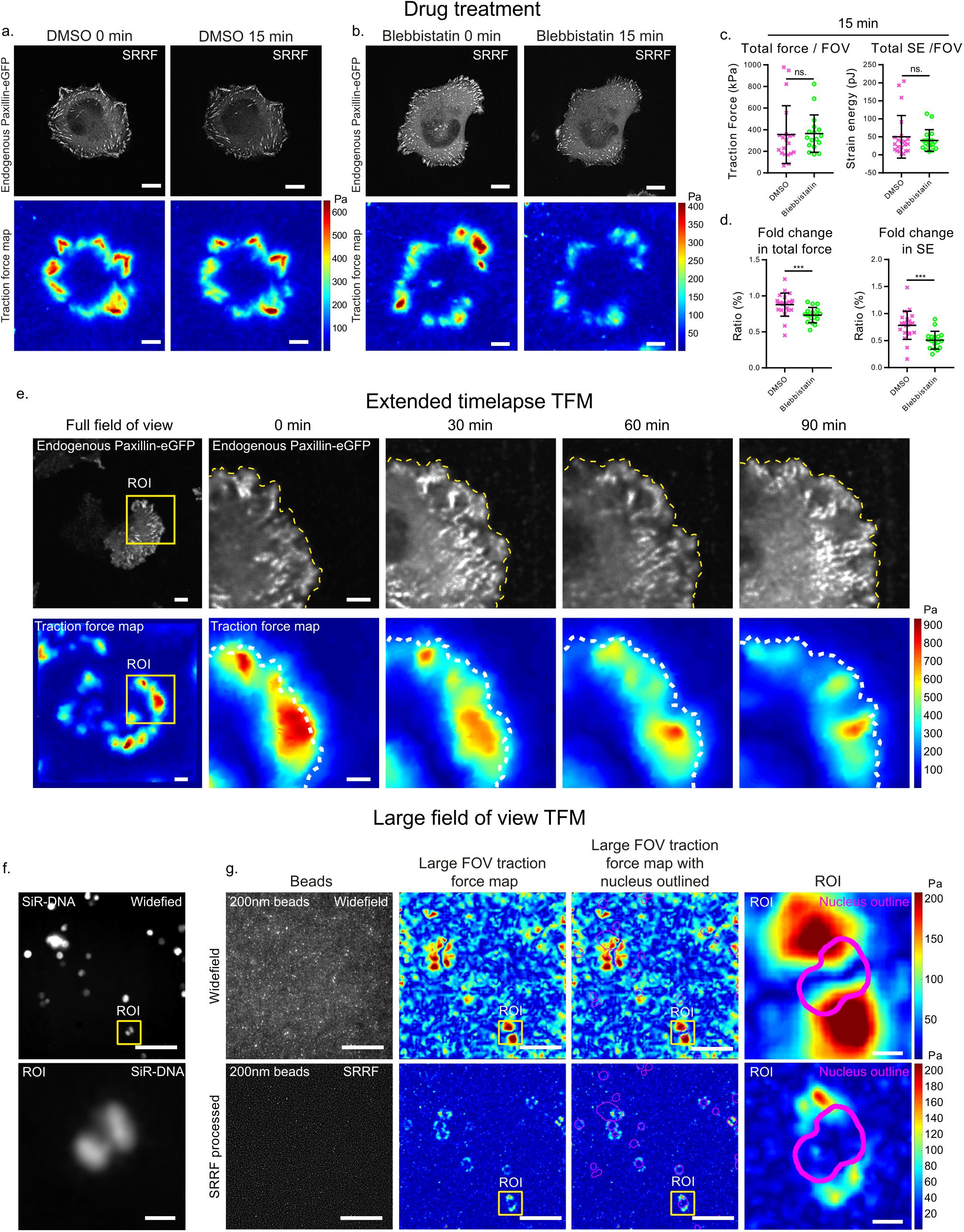
Applying FBSR-TFM to cell biological experiments. (**a,b**) U2OS cells expressing endogenously-tagged paxillin were plated on 9.6 kPa TFM gels containing 40 nm beads, and treated with either DMSO (**a**) or 10 µM blebbistatin for 15 min (**b**). Cells were imaged before and after treatment. FBSR was performed on both the bead and the paxillin images (LiveSRRF) and TFM analyses were performed using MATLAB. Representative images of cells and the corresponding traction maps are displayed. (**c,d**) Quantification of overall total forces and strain energy (SE) after treatments (**a,b**) (cropped to include only one cell) (**c**) and the fold change in total force and SE per field of view are displayed as dot plots (DMSO, n = 22; blebbistatin, n=17; 2 biological repeats). Statistics: Mann-Whitney U test. ***: p 0.004. (**e**) U-251 glioma cells expressing endogenously-tagged paxillin were plated on 9.6 kPa TFM gels containing 40 nm beads. Cells were imaged live, every 5 minutes and FBSR-TFM was performed (SDCM imaging, Live-SRRF processing and TFM analysis using MATLAB). The paxillin channel was denoised using the Noise2VOID algorithm (Krull et al., 2018). A representative field of view is displayed for the paxillin channel as well as the matching traction force map. The yellow square highlights a ROI that is magnified and displayed for several time points. The full movie is provided as supplemental information (Video 1). Dashed line depicts the leading edge of the cell. Scale bar: (main) 10 µm; (inset) 5 µm. (**f-g**) U2OS cells were plated on 2.6 kPa TFM gels containing 200 nm beads (classic protocol), treated with SiR-DNA to label nuclei (**f**), and imaged to perform FBSR TFM using a widefield microscope (20x air objective). The SiR-DNA images was denoised using the Noise2VOID algorithm (Krull et al., 2018) (**f**). Both widefield and FBSR images (LiveSRRF) were used to perform TFM analyses (MATLAB software). Images of the beads and the matching traction force maps are displayed (**g**). The outline of the nucleus is overlaid in magenta. Yellow squares highlight ROI that are magnified. Scale bar: (main) 100 µm; (inset) 10 µm.

FBSR is only mildly phototoxic, an important property for extended live-cell imaging (Culley et al., 2018b). Therefore, we next sought to assess if FBSR would be suitable for extended live TFM experiments. Glioma cells with endogenously tagged paxillin were imaged every 5 minutes, over a 100-minute time period and FBSR TFM measurements were performed (Figure 4e, Video 1 and Supplementary Figure 4a). In this experiment, the signal to noise ratio of endogenous paxillin was improved using a recent denoising approach based on convolutional neural network (Krull et al., 2018). Using these images, modulation of forces could clearly be observed, at high resolution, as cells protruded and migrated (Figure 4e, Video 1, Supplementary Figure 4a). Glioma cells imaged using this strategy were not visibly disturbed by the imaging. In addition, the same strategy could be used to perform extended live TFM imaging of human induced pluripotent stem cells, which, in our experience, are very sensitive to phototoxicity. Together this demonstrates that this method is suitable for long-term live TFM imaging (Supplementary Figure 4b).

One major Limitations of TFM is its low throughput, which considerably impedes the discovery of novel proteins / compounds that modulate force. While dedicated TFM screening platforms have been reported (Yoshie et al., 2018), performing TFM screens remain extremely challenging due to the requirement of high-resolution imaging to accurately measure bead displacement. Therefore, we tested the capability of FBSR to improve bead detections over very large fields of view (399 µm x 399 µm) using high numerical aperture low magnification objectives. In this case, cells were plated on gels containing 200 nm fluorescent beads and imaged using a 20x air objective on a widefield microscope. TFM analyses were performed on both the widefield and FBSR images (Figure 4f and 4g). Using the widefield images, the force maps obtained were of poor quality and relatively noisy with forces detected in cell-free areas (Figure 4g). In contrast, FBSR processing of the same field of view drastically improved the traction force maps, and the area of high forces closely matched the outline of the cells (Figure 4g). We believe that this last application of FBSR-TFM has the potential to considerably simplify the implementation of TFM screens and accelerate discoveries in mechanobiology as well as the identification of therapeutically valuable compounds / pathways.

## Discussion

Here we propose a simplified protocol and imaging strategy, relying on FBSR, which improve TFM measurements based on enhancements in both bead detection and accuracy of bead tracking. Importantly, FBSR-improved TFM data in combination with FBSR-enhanced staining of cellular proteins (e.g. paxillin) can be used to correlate force data with specific cellular structures such as focal adhesions. Our analysis pipeline can be adapted to camera-based confocal or widefield microscopes and is fully compatible with freely available TFM analysis software. In addition, we demonstrate that our workflow can be used to gain biologically relevant information and is suitable for long-term live measurement of traction forces. Finally, we propose that our strategy could be used to considerably simplify the implementation of TFM screens.

One current limitation of the FBSR-based TFM workflow described here is that it is currently not compatible with 3D TFM (3D tracking of beads underneath cells plated on a 2D substrate) as demonstrated recently using SIM (Colin-York et al., 2019). However, the strategy described here has not yet reached full potential and could be further developed. In particular, as FBSR reconstruction algorithms are under constant development and continue to improve (Dertinger et al., 2009; Gustafsson et al., 2016; Zhao et al., 2018),we expect parallel advances in the quality of FBSR TFM. For example, while FBSR algorithms can be capable of axial resolution improvement (Dertinger et al., 2009), this feature is not yet widely implemented and is likely to be improved in the future. In addition, as FBSR TFM is fully compatible with existing TFM software, it can be further developed by fine-tuning the computational algorithms responsible for bead recognition, tracking and methods used to derive cellular forces (Han et al., 2015). Here, we principally used the FTTC method to reconstruct forces, but it is tempting to speculate that even further quality enhancement could be gained by employing more computationally heavy mathematical frameworks (Han et al., 2015). In addition, it is theoretically possible that the resolution of FBSR TFM could be further enriched by mixing beads of different colors as demonstrated for confocal-based microscopy (Plotnikov et al., 2014).

## Materials and methods

### Cell culture and transient transfection

U2OS and U-251 glioma cells were grown in DMEM/F-12 (Dulbecco’s Modified Eagle’s Medium/Nutrient Mixture F-12; Life Technologies, 10565-018) supplemented with 10% fetal bovine serum (FCS) (Biowest, S1860). U2OS cells were purchased from DSMZ (Leibniz Institute DSMZ-German Collection of Microorganisms and Cell Cultures, Braunschweig DE, ACC 785). U-251 glioma cells were a generous gift from Professor David Odde (University of Minnesota, US). The human induced pluripotent stem cell (hPSC) line HEL24.3 was a kind gift from Professor Timo Otonkoski (University of Helsinki). This cell line was created using Sendai viruses (Trokovic et al., 2015) and was cultured on Matrigel (Corning, 354277) in Essential 8 Basal medium (Life Technologies, A15169-01) supplemented with E8 supplements (Life Technologies, A1517-01). Cells expressing endogenously tagged paxillin-GFP were generated using CRISPR / Cas9 (Roberts et al., 2017). The gRNA sequence targeting paxillin (5’-GCACCTAGCAGAAGAGCTTG-3’) was cloned into the pSpCas9(BB)-2A-GFP backbone using the BbsI restriction site (Roberts et al., 2017). Cells were then transfected with the GFP-Cas9-paxillin_gRNA construct and the template plasmid AICSDP-1:PXN-EGFP in equimolar ratio (1:1). After transfection, cells were grown for 5 days before being sorted based on green fluorescence using a fluorescence-activated cell sorter (FACS; FACSAria IIu, BD). AICSDP-1:PXN-EGFP was a gift from The Allen Institute for Cell Science (Addgene plasmid 87420).

### TFM gel preparation

35 mm glass bottom dishes (Cellvis, D35-14-1N) were treated with Bind-Silane solution (GE Healthcare, Silane A-174) for 15 minutes at RT, washed once with 95 % EtOH and twice with mQH2O before being left to dry completely. In parallel, 13 mm glass coverslips were coated with Poly-D-lysine (Sigma-Aldrich, A-003-E) for 20 min at +4°C, washed in mQH2O and then left to dry out. Fluorescent beads (either dark red, excitation 660 nm / emission 680 nm; or orange, excitation 540 nm / emission 560 nm; Thermo Fisher Scientific, F10720) were diluted 1:5000 in mQH2O. The bead solution was then subjected to repeated sonication for 30 seconds, followed by a 30-second pause, over a 10-minute period. Importantly, beads were kept on ice for this entire duration to prevent bead clustering. Each Poly-D-lysine coated coverslip was then incubated with a 150 µl drop of the bead solution at +4°C for 20 min. Coverslip were then washed and kept in mQH2O. Before their use, the glass coverslips were left to dry completely.

A pre-mixture composed of 40% acrylamide (Sigma-Aldrich, A4058), 2% N, N-Methylenebisacrylamide solution (Sigma-Aldrich, M1533) and PBS was prepared according to the desired gel stiffness (see Table 2). In this study, most experiments were performed using ~ 9.6 kPa TFM gels with the exception of hPSC live imaging (~ 32 kPa gel; Supplementary Figure 4b) and of the large field of view TFM (~2.6 kPa gel, Figure 4f and 4g).

**Table 2.** TFM gel pre-mixture recipe.

| AA. 40% (μl) | Bis. AA. 2% μl | PBS (μl) | 200 nm beads (ul), optional | APS. 10% (μl) | TEMED (μl) | E (kPa) |
| --- | --- | --- | --- | --- | --- | --- |
| 94 | 15 | 346 | 3.4 | 5 | 1 | ~2.6 |
| 94 | 50 | 356 | 3.4 | 5 | 1 | ~9.6 |
| 225 | 100 | 175 | 3.4 | 5 | 1 | ~31.7 |

From this stage onward, the pre-mixture was then kept on ice and sonicated for 30 seconds followed by a 30 second pause over 10 minutes. The pre-mixture was then vortexed briefly, and 0.2% TEMED (vol/vol; Sigma-Aldrich, T9281) and 1% ammonium persulfate (vol/vol; 10% stock solution) were added to start the PAA polymerization. After a brief vortex, 11.8 µl of gel mixture was added onto the glass of each glass bottom dish. The bead-coated 13 mm coverslip was then carefully placed on top of the drop, ensuring that a thin layer of liquid remained between the two glass surfaces. The plates were then incubated at room temperature for 60 min after which they were submerged in PBS to allow the careful removal of the top glass coverslip using forceps.

In the experiments where SDCM TFM was performed in addition to FBSR TFM, 200nm green fluorescent beads (excitation 505 nm / emission 515 nm) (Life Technologies, F8811) were also added to the pre-mixture prior to the sonication step (Table 2).

To allow for functionalization, the TFM gels were incubated for 30 minutes at RT with an activation solution [0.2 mg/ml of Sulfo-SANPAH (Thermo Fisher Scientific, 22589), 2 mg/ml N-(3-Dimethylaminopropyl)-N-ethylcarbodiimide hydrochloride (EDC) (Sigma-Aldrich, 03450) diluted in 50 mM HEPES (4-(2-hydroxyethyl)-1-piperazineethanesulfonic acid; Sigma-Aldrich, H0887) under gentle agitation. Glass bottom dishes containing the gels were then irradiated, without their plastic lids, with ultraviolet (UV) light for 10 minutes using a UV-chamber (Jelight Company Inc., UVO CLEANER, 342-220). Gels were then washed three times with PBS and coated with either 10 µg/ml fibronectin (Merck-Millipore, 341631) (U2OS and U-251 cells) or 5 µg/ml vitronectin (Thermo Fisher Scientific, A14700) (hPSCs).

### Microcopy setup

The spinning disk confocal microscope (SDCM) used was a Marianas spinning disk imaging system with a Yokogawa CSU-W1 scanning unit on an inverted Zeiss Axio Observer Z1 microscope controlled by SlideBook 6 (Intelligent Imaging Innovations, Inc.). Images were acquired using a Photometrics Evolve, Back Illuminated EMCCD camera (512 x 512 pixels). The microscope was used either in confocal or widefield mode as indicated in the figure legends. Objectives used were a 20x (NA 0.8 air, Plan Apochromat, DIC) objective (Zeiss) and a 63x (NA 1.15 water, LD C-Apochromat) objective (Zeiss).

### Traction force microscopy

Cells were plated on TFM gels (in a 1 ml volume of media) and left to adhere for at least 2 hours prior to imaging. To avoid drifting during imaging, the spinning disk microscope was pre-warmed to 37°C prior to image acquisition. To perform TFM measurements, beads were imaged before (Pre) and after (Post) removing cells, the Pre and Post images were then aligned (see methods below), the beads detected in both images and their movements tracked and local forces measured (see methods below). To perform FBSR, 100 frames of the Pre and Post bead planes and of the paxillin staining (when required) were acquired. In between acquiring the Pre and Post images, the cells were removed by adding 500 µl of a 20% SDS solution (5-minute incubation).

To perform the blebbistatin treatment experiment (Figure 4a), a first set of FBSR images of beads and focal adhesions were acquired (first Pre image). Then a warm solution containing 20 µM blebbistatin (final concentration in cell medium; Stemcell Technologies, 72402) or DMSO (Sigma-Aldrich, D2650) was added to the cells (1 ml added) for 15 minutes. A second set of FBSR images of the beads and focal adhesions was then acquired (second Pre image). The cells were then detached as described above and the final set of FBSR images was then acquired (Post image). To perform extended live TFM imaging of the iPSCs and U-251 cells, FBSR image sets of the beads were acquired every 5 minutes and the cells were detached as described above.

### Image alignment

Prior to bead tracking and force mapping, the Pre and Post bead images were aligned in the Fiji distribution of ImageJ (Schindelin et al., 2012; Rueden et al., 2017; Schneider et al., 2012) using either the NanoJ-Core (Laine et al., 2019) or the “Linear stack alignment with SIFT” plugins. The “Linear stack alignment with SIFT” plugin was only used when affine registration was required. In all cases, the first Pre image was used as a reference image. For the blebbistatin and for the live TFM experiments, the focal adhesion images were registered with the bead images using the drift table generated by the NanoJ-Core plugin (Laine et al., 2019).

### FBSR processing

FBSR processing was performed using NanoJ-LiveSRRF and SACD (Zhao et al., 2018). NanoJ-LiveSRRF is the newest implementation of NanoJ-SRRF within the ImageJ software. NanoJ-LiveSRRF is available upon request (will be openly available for download soon), whereas NanoJ-SRRF is already an open source software (Gustafsson et al., 2016).

For FBSR processing using LiveSRRF, 100 frames were used for the reconstruction. The parameter sweep option as well as SQUIRREL analyses (RSE and RSP values) (Culley et al., 2018a), integrated within LiveSRRF, were used to define the optimal reconstruction parameters (32 different conditions as shown in Supplementary Figure 1b). For each liveSRRF setting combination, RSP and RSE values, and bead density were measured and ranked. The optimal LiveSRRF settings were then determined based on all criteria (best overall rank) (see Supplementary Figure 1c) and the final parameters used to process images are listed in table 3.

**Table 3.**
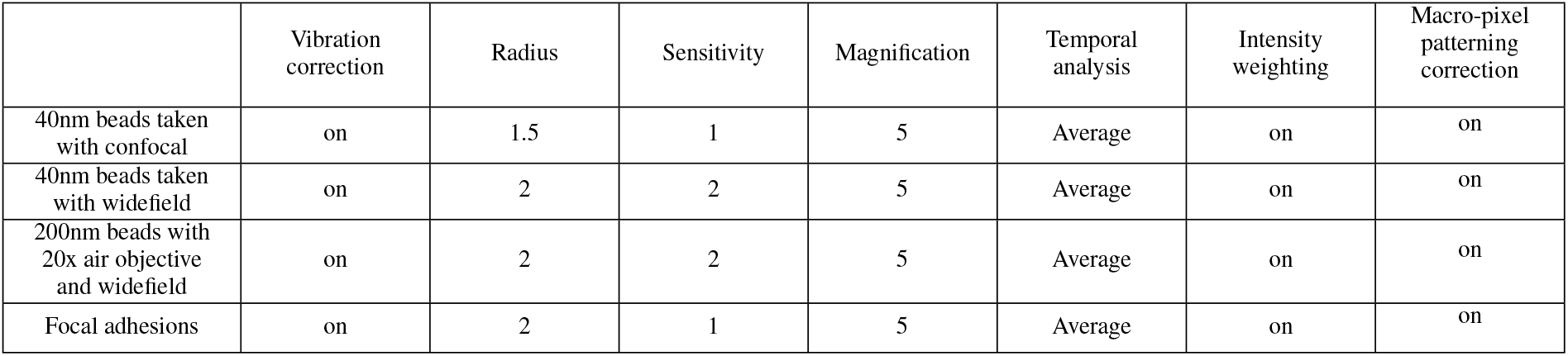
LiveSRRF parameters.

|  | Vibration correction | Radius | Sensitivity | Magnification | Temporal analysis | Intensity weighting | Macro-pixel patterning correction |
| --- | --- | --- | --- | --- | --- | --- | --- |
| 40nm beads taken with confocal | on | 1.5 | 1 | 5 | Average | on | on |
| 40nm beads taken with widefield | on | 2 | 2 | 5 | Average | on | on |
| 200nm beads with 20x air objective and widefield | on | 2 | 2 | 5 | Average | on | on |
| Focal adhesions | on | 2 | 1 | 5 | Average | on | on |

For FBSR processing with SACD (Zhao et al., 2018), the first 50 frames were used for the reconstruction. SACD reconstructions were performed within MATLAB (Mathworks, version R2019a) using the following parameters (NA, 1.15; pixel size, 247 × 10^*−*9^; lamda, 647 × 10^*−*9^; iter, 1; mag, 5; squre, 2, order, 3). The MATLAB script used to process SACD images is available on GitHub and can be found at https://github.com/guijacquemet/.

### Quantification of bead density

Bead density was quantified by dividing the number of beads detected (in a field of view) by the size of the field of view. The number of beads for each field of view was measured in Fiji using the “find maxima” option. The threshold used was tuned for each dataset so that only beads were counted. Importantly, the number of beads measured using this strategy was nearly identical to the number of beads identified by the MATLAB-based TFM software.

### Bead tracking and local force measurements

The bead tracking and local force measurements were performed either using MATLAB (Mathworks, version R2019a) or using Fiji (Schindelin et al., 2012; Rueden et al., 2017; Schneider et al., 2012). For the MATLAB-based analyses, the TFM software developed by the Danuser laboratory was used (Han et al., 2015). Key parameters used can be found in table 4.

**Table 4.** Key parameters used to perform TFM analyses using the MATLAB-based software

|  | Bead detection parameters | “Template size” and “maximum displacement for calculating displacement field” | Force field calculation |
| --- | --- | --- | --- |
| Figure 2 | High-resolution subsampling of beads and Use subpixel correlation via image interpolation | 20 and 21 px | FTTC (Fourier-Transform Traction Cytometry) |
| Figure 3 | High-resolution subsampling of beads and Use subpixel correlation via image interpolation | 40 and 41 px | FTTC |
| Figure 4a | High-resolution subsampling of beads and Use subpixel correlation via image interpolation | 80 and 81 px | FTTC |
| Figure 4e | PIV | 80 and 81 px | FTTC |
| Figure 4g | High-resolution subsampling of beads and Use subpixel correlation via image interpolation | 80 and 81 px | FTTC |

To generate the displacement and traction maps in Fiji, the particle image velocity (PIV) plugin and the Fourier transformation traction cytometry (FTTC) plugin (Tseng et al., 2012) were used. The aligned images of the Pre- and Post TFM images of the beads were first processed with the PIV plugin using the correlation coefficient iteration option (interrogation window sizes: 128 pixel first round, 64 pixel second round and 32 pixel third round). The resulting PIV text file was then saved and plotted as a displacement map using the plot function. Images shown in figure 3c were post-processed using the normalized median test option (parameters used: 0.2 for noise and 5.0 for threshold). The traction force maps were generated using the ImageJ FTTC plugin (poisson ratio, 0.5; Young’s modulus; 10 kPa; regularisation parameter, 4.0 × 10^*−*10^). The total forces were calculated by measuring the integrated density of the 32-bit images produced by the plugin.

### Image denoising using Noise2VOID

The signal to noise ratio of endogenously tagged paxillin (Figure 4e) and of DNA (Sir-DNA) (Figure 4f) were improved using the recent denoising approach Noise2VOID (Krull et al., 2018) which is based on convolutional neural networks. Noise2VOID was executed through the Google co-laboratory platform, which can run Jupyter Notebooks in the cloud. The Jupyter Notebooks used are available on GitHub and can be found at https://github.com/guijacquemet/.

### Statistical analysis

Correlation analyses were performed using Spearman’s Rank-order. Statistical analyses were performed using the Mann-Whitney U test. The error bars presented in figures depict the standard deviation. N numbers are indicated in the figure legends.

### Data availability

The authors declare that the data supporting the findings of this study are available within the article and from the authors on request.

## Acknowledgements

We thank Aleksi Isomursu for providing us the U-251 glioma cell line expressing endogenously-tagged paxillin-GFP. We thank Dr Hellyeh Hamidi for critical reading of the manuscript. We thank J. Siivonen and P. Laasola for technical assistance and M. Saari for help with the microscopes. The Cell Imaging and Cytometry core facility (Turku Bioscience, University of Turku, Åbo Akademi University and Biocenter Finland) is acknowledged for services, instrumentation, and expertise. The Allen Institute for Cell Science, Professor David Odde (University of Minnesota, US) and Professor Timo Otonkoski (University of Helsinki) are acknowledged for providing reagents.

This study has been supported by the Academy of Finland (G.J., and J.I.), the Academy of Finland CoE for Translational Cancer Research (J.I.), an ERC CoG grant (615258), the Sigrid Juselius Foundation and the Finnish Cancer Organization (J.I. and G.J.). A.S. has been supported by the University of Turku Doctoral programme for Molecular Medicine (TuDMM). R.F.L and R.H were funded by grants from the UK Biotechnology and Biological Sciences Research Council [BB/S507532/1], the UK Medical Research Council [MR/K015826/1], the Wellcome Trust [203276/Z/16/Z] and core funding by the MRC Laboratory for Molecular Cell Biology, University College London [MC_UU12018/7].

## Authors contributions

Conceptualization, A.S., G.J. and J.I.; Methodology, A.S., G.J.; Formal Analysis, G.J., C.G. and A.S.; Investigation, A.S.; Resources, R.L. and R.H.,; Writing – Original Draft, A.S., G.J. and J.I.; Writing – Review and Editing, A.S., R.L., C.G., R.H, G.J. and J.I.; Visualization, A.S., G.J. and J.I.; Supervision, G.J. and J.I.; Funding Acquisition, G.J. and J.I.

## Supplementary material

### Video 1. Live FBSR TFM in a migrating glioma cell

U-251 glioma cells expressing endogenously-tagged paxillin were plated on 9.6 kPa TFM gels containing 40 nm beads. Cells were imaged live, every 5 minutes, using a spinning disk microscope to allow FBSR processing of the beads (LiveSRRF) and subsequent TFM analysis using the MATLAB-based software. The paxillin channel (denoised using Noise2VOID) as well as the matching traction force and bead displacement maps are displayed. The yellow square highlights a ROI that is magnified. Scale bars 10 µm.

---

**Figure S1:**
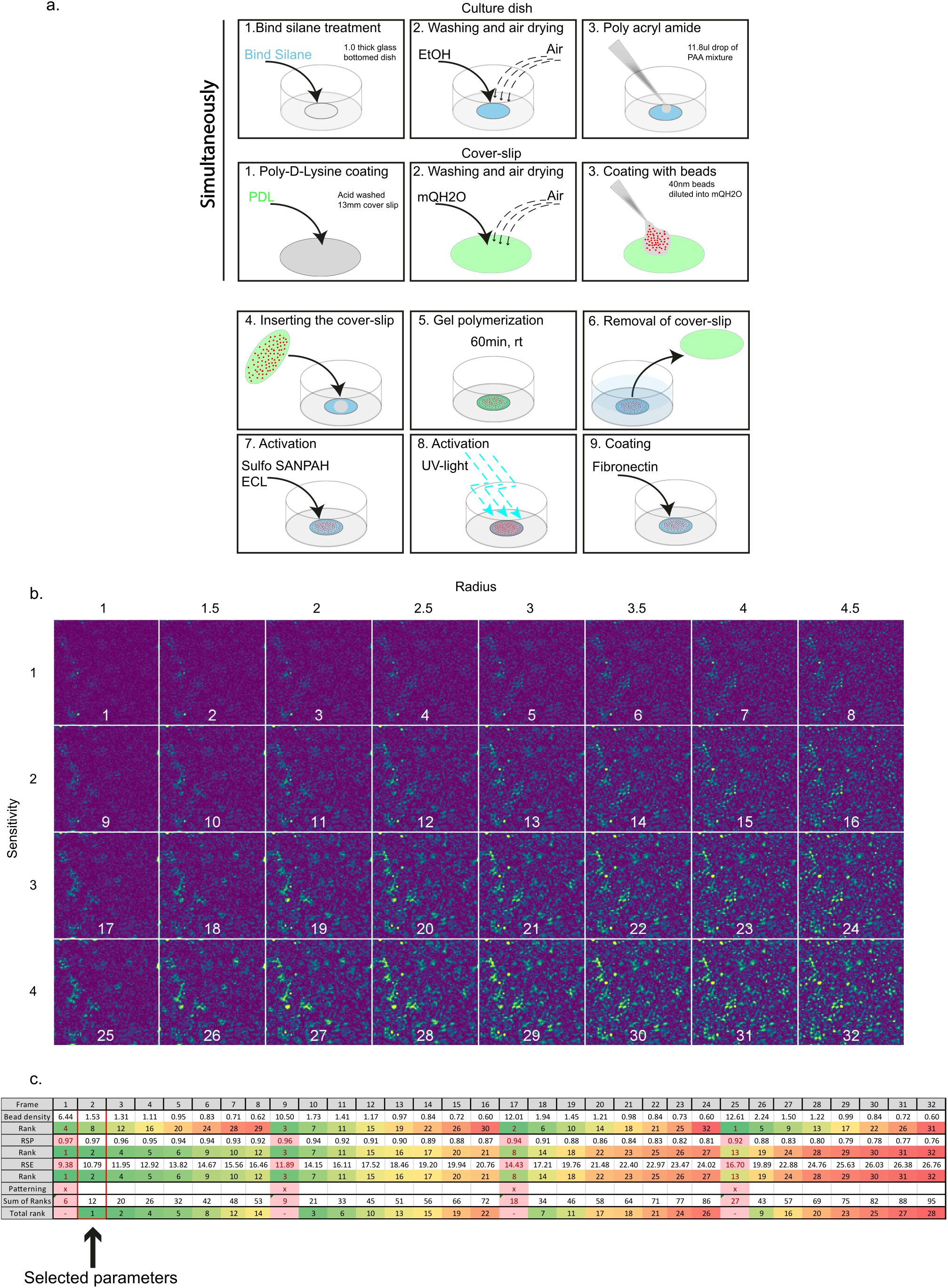
Beads detection within TFM gels using FBSR. (**a**) A schematic cartoon illustrating the protocol used to generate TFM gels with 40 nm beads embedded only on the top layer. (**b-c**) To determine the optimal LiveSRRF settings to improve bead images, the field of view displayed in figure 1c was processed with 32 different combinations of settings (by changing the radius and sensitivity values as indicated) and the resulting images were analysed using NanoJ-SQUIRREL. The resulting error maps are displayed as a montage (**b**) and are color-coded to visualise areas of low (purple) and high error (yellow). Each combination was then ranked (from 1 (green) to 32 (red)) according, RSP and RSE values and bead density (**c**). The overall best ranked combination was chosen as the final settings for optimal LiveSRRF processing.

**Figure S2:**
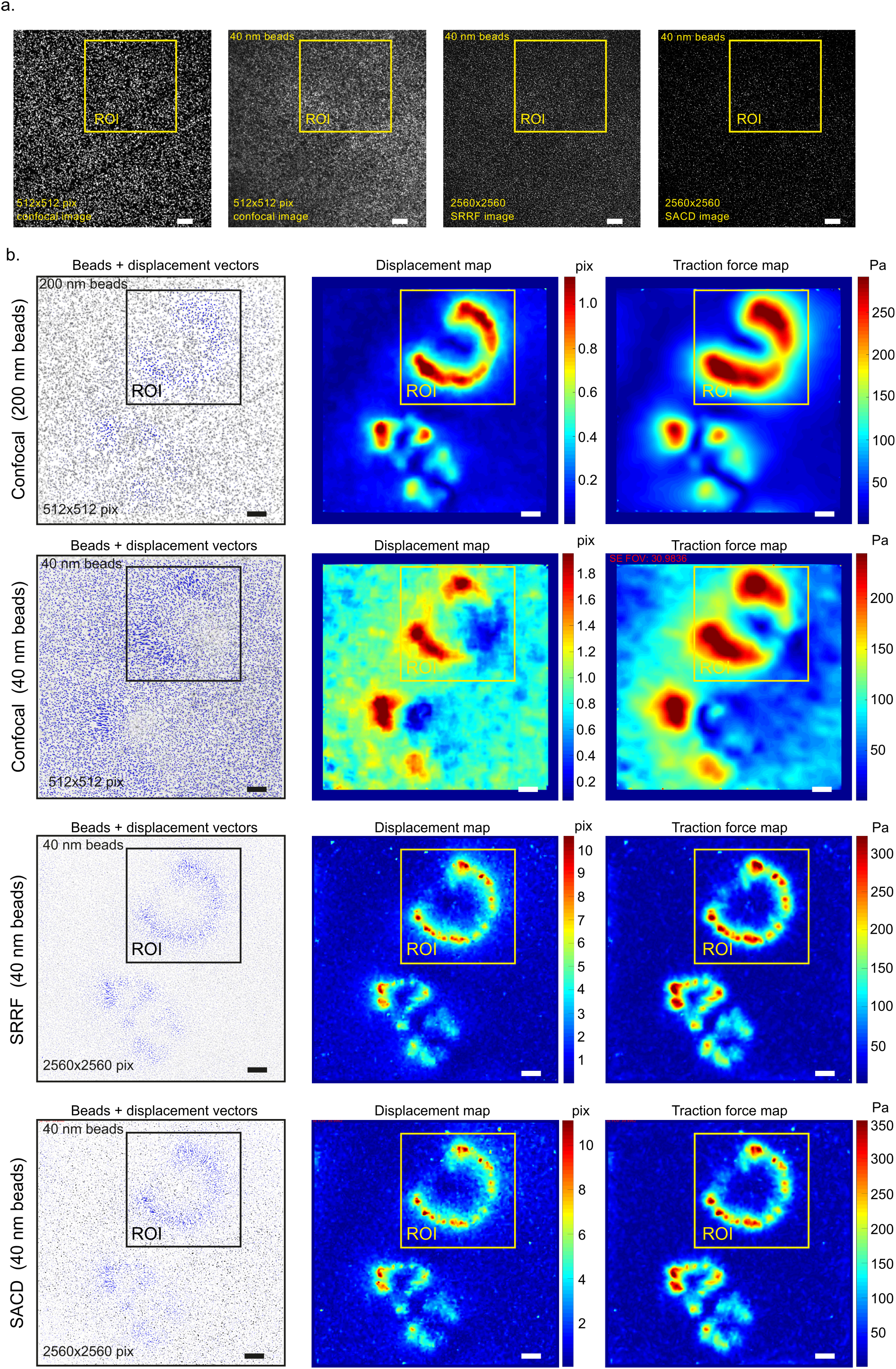
Full field of view for figure 2. Full field of view of the images and maps presented in figure 2. The yellow and black squares highlight the ROI used to create figure 2. Scale bar 10 µm.

**Figure S3:**
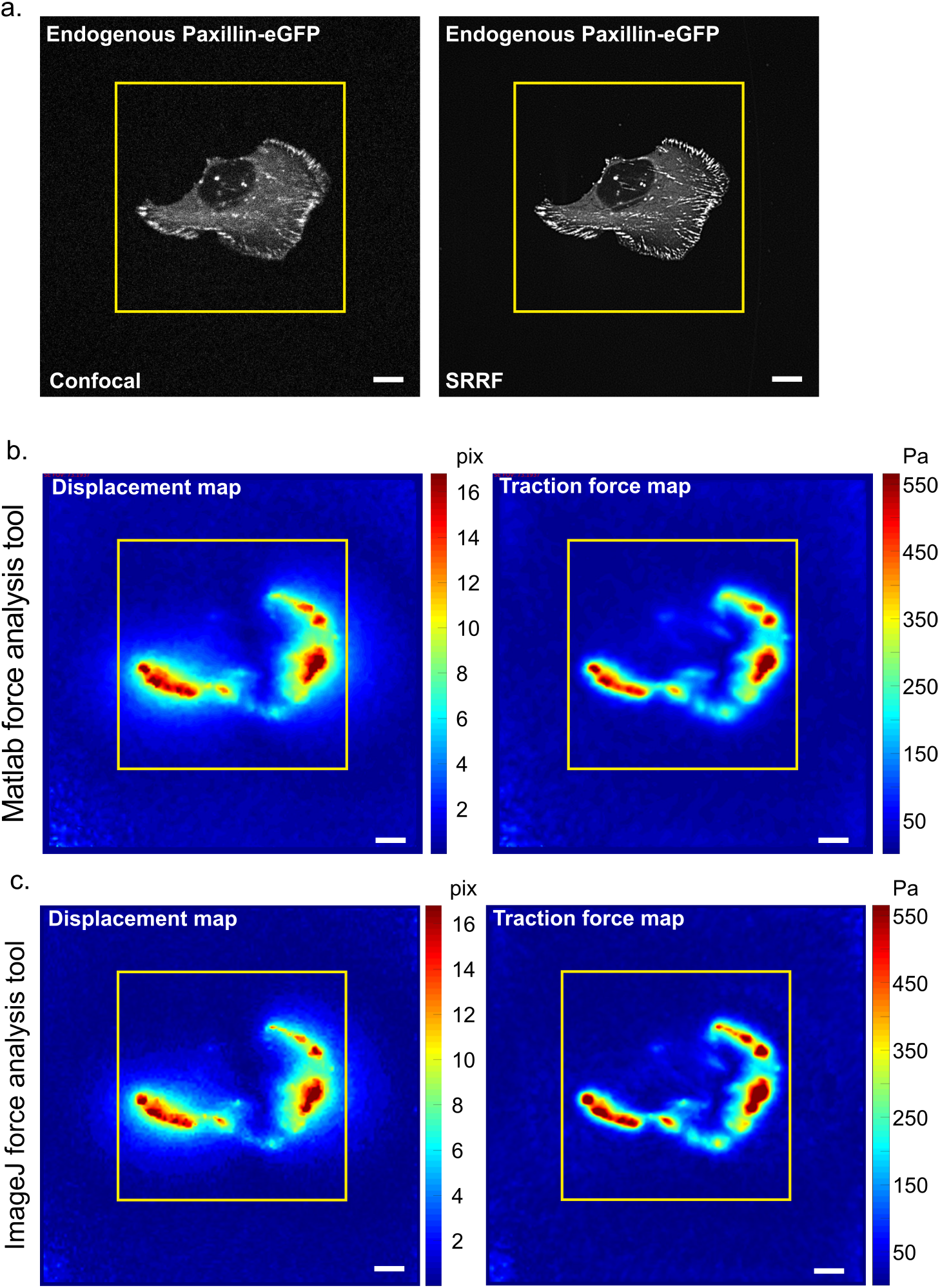
Full field of view for figure 3. Full field of view of the images and maps presented in figure 3. The yellow squares highlight the ROI used to create figure 3 Scale bar 10 µm.

**Figure S4:**
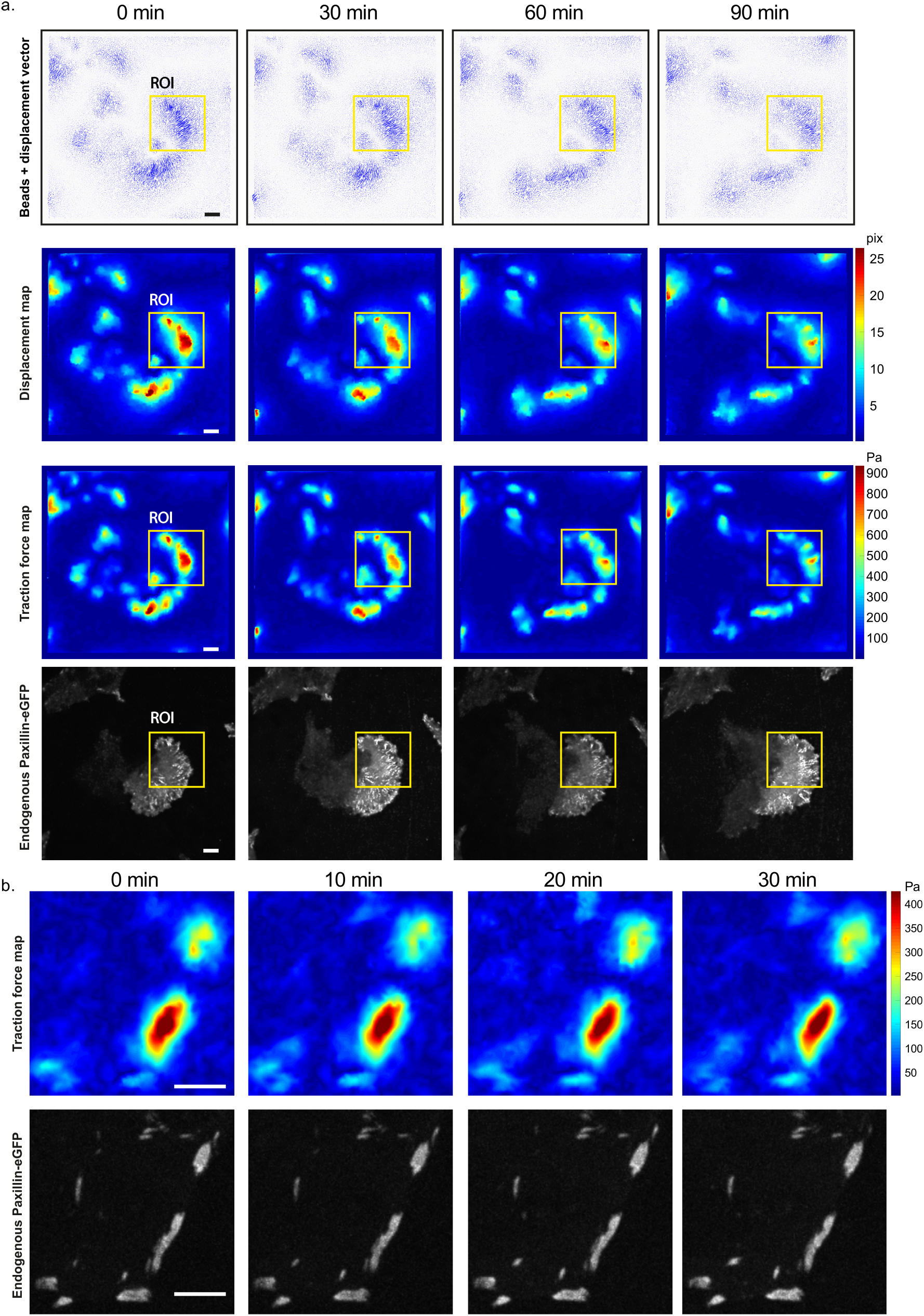
Implementation of fluctuation-based TFM for extended live cell imaging. (**a**) Full field of view of the images and maps presented in figure 4e. In addition, the bead displacement maps as well as images of the beads (black) with the bead movement indicated by displacement vectors (purple arrows, length scaled up by 2) are displayed. The yellow squares indicate the ROI used to create figure 4e. Scale bars 10 µm. (**b**) Human induced pluripotent stem cells expressing endogenously-tagged paxillin were plated on 30 kPa TFM gels containing 40 nm beads. Cells were imaged live, every 5 minutes, using a spinning disk microscope to allow the FBSR processing of the beads and subsequent TFM analysis using the MATLAB-based software. A representative ROI for the paxillin channel and the corresponding traction force map are displayed for the different time points.

## Bibliography

Bagshaw, C.R., and D. Cherny. 2006. Blinking fluorophores: what do they tell us about protein dynamics? Biochem. Soc. Trans. 34:979–982. doi:10.1042/BST0340979.

Broders-Bondon, F., T.H.N. Ho-Bouldoires, M.-E. Fernandez-Sanchez, and E. Farge. 2018. Mechanotransduction in tumor progression: The dark side of the force. J. Cell Biol. 217:1571–1587. doi:10.1083/jcb.201701039.

Colin-York, H., Y. Javanmardi, L. Barbieri, D. Li, K. Korobchevskaya, Y. Guo, C. Hall, A. Taylor, S. Khuon, G.K. Sheridan, T.-L. Chew, D. Li, E. Moeendarbary, and M. Fritzsche. 2019. Spatiotemporally Super-Resolved Volumetric Traction Force Microscopy. Nano Lett. 19:4427–4434. doi:10.1021/acs.nanolett.9b01196.

Colin-York, H., D. Shrestha, J.H. Felce, D. Waithe, E. Moeendarbary, S.J. Davis, C. Eggeling, and M. Fritzsche. 2016. Super-Resolved Traction Force Microscopy (STFM). Nano Lett. 16:2633–2638. doi:10.1021/acs.nanolett.6b00273.

Conway, J.R.W., and G. Jacquemet. 2019. Cell matrix adhesion in cell migration. Essays Biochem. EBC20190012. doi:10.1042/EBC20190012.

Culley, S., D. Albrecht, C. Jacobs, P.M. Pereira, C. Leterrier, J. Mercer, and R. Henriques. 2018a. Quantitative mapping and minimization of super-resolution optical imaging artifacts. Nat. Methods. 15:263–266. doi:10.1038/nmeth.4605.

Culley, S., K.L. Tosheva, P. Matos Pereira, and R. Henriques. 2018b. SRRF: Universal live-cell super-resolution microscopy. Int. J. Biochem. Cell Biol. 101:74–79. doi:10.1016/j.biocel.2018.05.014.

Dertinger, T., R. Colyer, G. Iyer, S. Weiss, and J. Enderlein. 2009. Fast, background-free, 3D super-resolution optical fluctuation imaging (SOFI). Proc. Natl. Acad. Sci. 106:22287–22292. doi:10.1073/pnas.0907866106.

Duscher, D., Z.N. Maan, V.W. Wong, R.C. Rennert, M. Januszyk, M. Rodrigues, M. Hu, A.J. Whitmore, A.J. Whittam, M.T. Longaker, and G.C. Gurtner. 2014. Mechanotransduction and fibrosis. J. Biomech. 47:1997–2005. doi:10.1016/j.jbiomech.2014.03.031.

Georgiadou, M., J. Lilja, G. Jacquemet, C. Guzmán, M. Rafaeva, C. Alibert, Y. Yan, P. Sahgal, M. Lerche, J.-B. Manneville, T.P. Mäkelä, and J. Ivaska. 2017. AMPK negatively regulates tensin-dependent integrin activity. J. Cell Biol. 216:1107–1121. doi:10.1083/jcb.201609066.

Gustafsson, N., S. Culley, G. Ashdown, D.M. Owen, P.M. Pereira, and R. Henriques. 2016. Fast live-cell conventional fluorophore nanoscopy with ImageJ through superresolution radial fluctuations. Nat. Commun. 7:12471. doi:10.1038/ncomms12471.

Hakanpaa, L., E.A. Kiss, G. Jacquemet, I. Miinalainen, M. Lerche, C. Guzmán, E. Mervaala, L. Eklund, J. Ivaska, and P. Saharinen. 2018. Targeting β1-integrin inhibits vascular leakage in endotoxemia. Proc. Natl. Acad. Sci. U. S. A. 115:E6467–E6476. doi:10.1073/pnas.1722317115.

Han, S.J., Y. Oak, A. Groisman, and G. Danuser. 2015. Traction microscopy to identify force modulation in subresolution adhesions. Nat. Methods. 12:653–656. doi:10.1038/nmeth.3430.

Heisenberg, C.-P., and Y. Bellaïche. 2013. Forces in Tissue Morphogenesis and Patterning. Cell. 153:948–962. doi:10.1016/j.cell.2013.05.008.

Jacquemet, G., A. Stubb, R. Saup, M. Miihkinen, E. Kremneva, H. Hamidi, and J. Ivaska. 2019. Filopodome Mapping Identifies p130Cas as a Mechanosensitive Regulator of Filopodia Stability. Curr. Biol. 29:202–216.e7. doi:10.1016/j.cub.2018.11.053.

Kechagia, J.Z., J. Ivaska, and P. Roca-Cusachs. 2019. Integrins as biomechanical sensors of the microenvironment. Nat. Rev. Mol. Cell Biol. 20:457–473. doi:10.1038/s41580-019-0134-2.

Krull, A., T.-O. Buchholz, and F. Jug. 2018. Noise2Void - Learning Denoising from Single Noisy Images. ArXiv181110980 Cs.

Laine, R.F., K.L. Tosheva, N. Gustafsson, R.D.M. Gray, P. Almada, D. Albrecht, G.T. Risa, F. Hurtig, A.-C.L. as, B. Baum, J. Mercer, C. Leterrier, P.M. Pereira, S. Culley, and R. Henriques. 2019. NanoJ: a high-performance open-source super-resolution microscopy toolbox. J. Phys. Appl. Phys. 52:163001. doi:10.1088/1361-6463/ab0261.

Lampi, M.C., and C.A. Reinhart-King. 2018. Targeting extracellular matrix stiffness to attenuate disease: From molecular mechanisms to clinical trials. Sci. Transl. Med. 10. doi:10.1126/scitranslmed.aao0475.

Linde, S. van de, and M. Sauer. 2014. How to switch a fluorophore: from undesired blinking to controlled photoswitching. Chem. Soc. Rev. 43:1076–1087. doi:10.1039/C3CS60195A.

Närvä, E., A. Stubb, C. Guzmán, M. Blomqvist, D. Balboa, M. Lerche, M. Saari, T. Otonkoski, and J. Ivaska. 2017. A Strong Contractile Actin Fence and Large Adhesions Direct Human Pluripotent Colony Morphology and Adhesion. Stem Cell Rep. 9:67–76. doi:10.1016/j.stemcr.2017.05.021.

Plotnikov, S.V., B. Sabass, U.S. Schwarz, and C.M. Waterman. 2014. Chapter 20 - High-Resolution Traction Force Microscopy. In Methods in Cell Biology. J.C. Waters and T. Wittman, editors. Academic Press. 367–394.

Roberts, B., A. Haupt, A. Tucker, T. Grancharova, J. Arakaki, M.A. Fuqua, A. Nelson, C. Hookway, S.A. Ludmann, I.A. Mueller, R. Yang, R. Horwitz, S.M. Rafelski, and R.N. Gunawardane. 2017. Systematic gene tagging using CRISPR/Cas9 in human stem cells to illuminate cell organization. Mol. Biol. Cell. 28:2854–2874. doi:10.1091/mbc.E17-03-0209.

Roca-Cusachs, P., V. Conte, and X. Trepat. 2017. Quantifying forces in cell biology. Nat. Cell Biol. 19:742–751. doi:10.1038/ncb3564.

Rueden, C.T., J. Schindelin, M.C. Hiner, B.E. DeZonia, A.E. Walter, E.T. Arena, and K.W. Eliceiri. 2017. ImageJ2: ImageJ for the next generation of scientific image data. BMC Bioinformatics. 18:529. doi:10.1186/s12859-017-1934-z.

Schindelin, J., I. Arganda-Carreras, E. Frise, V. Kaynig, M. Longair, T. Pietzsch, S. Preibisch, C. Rueden, S. Saalfeld, B. Schmid, J.-Y. Tinevez, D.J. White, V. Hartenstein, K. Eliceiri, P. Tomancak, and A. Cardona. 2012. Fiji: an open-source platform for biological-image analysis. Nat. Methods. 9:676–682. doi:10.1038/nmeth.2019.

Schneider, C.A., W.S. Rasband, and K.W. Eliceiri. 2012. NIH Image to ImageJ: 25 years of image analysis. Nat. Methods. 9:671–675. doi:10.1038/nmeth.2089.

Segel, M., B. Neumann, M.F.E. Hill, I.P. Weber, C. Viscomi, C. Zhao, A. Young, C.C. Agley, A.J. Thompson, G.A. Gonzalez, A. Sharma, S. Holmqvist, D.H. Rowitch, K. Franze, R.J.M. Franklin, and K.J. Chalut. 2019. Niche stiffness underlies the ageing of central nervous system progenitor cells. Nature. 573:130–134. doi:10.1038/s41586-019-1484-9.

Style, R.W., R. Boltyanskiy, G.K. German, C. Hyland, C.W. MacMinn, A.F. Mertz, L.A. Wilen, Y. Xu, and E.R. Dufresne. 2014. Traction force microscopy in physics and biology. Soft Matter. 10:4047–4055. doi:10.1039/C4SM00264D.

Trokovic, R., J. Weltner, P. Noisa, T. Raivio, and T. Otonkoski. 2015. Combined negative effect of donor age and time in culture on the reprogramming efficiency into induced pluripotent stem cells. Stem Cell Res. 15:254–262. doi:10.1016/j.scr.2015.06.001.

Tseng, Q., E. Duchemin-Pelletier, A. Deshiere, M. Balland, H. Guillou, O. Filhol, and M. Théry. 2012. Spatial organization of the extracellular matrix regulates cell–cell junction positioning. Proc. Natl. Acad. Sci. 201106377. doi:10.1073/pnas.1106377109.

Wickström, S.A., and C.M. Niessen. 2018. Cell adhesion and mechanics as drivers of tissue organization and differentiation: local cues for large scale organization. Curr. Opin. Cell Biol. 54:89–97. doi:10.1016/j.ceb.2018.05.003.

Yoshie, H., N. Koushki, R. Kaviani, M. Tabatabaei, K. Rajendran, Q. Dang, A. Husain, S. Yao, C. Li, J.K. Sullivan, M. Saint-Geniez, R. Krishnan, and A.J. Ehrlicher. 2018. Traction Force Screening Enabled by Compliant PDMS Elastomers. Biophys. J. 114:2194–2199. doi:10.1016/j.bpj.2018.02.045.

Zhao, W., J. Liu, C. Kong, Y. Zhao, C. Guo, C. Liu, X. Ding, X. Ding, J. Tan, and H. Li. 2018. Faster super-resolution imaging with auto-correlation two-step deconvolution. ArXiv180907410 Phys.

Zündel, M., A.E. Ehret, and E. Mazza. 2017. Factors influencing the determination of cell traction forces. PLoS ONE. 12. doi:10.1371/journal.pone.0172927.

